# Categorical encoding of decision variables in orbitofrontal cortex

**DOI:** 10.1101/479519

**Authors:** Arno Onken, Jue Xie, Stefano Panzeri, Camillo Padoa-Schioppa

## Abstract

A fundamental and recurrent question in systems neuroscience is that of assessing what variables are encoded by a given population of neurons. Such assessments are often challenging because neurons in one brain area may encode multiple variables, and because neuronal representations might be categorical (different neurons encoding different variables) or mixed (individual neurons encoding a combination of variables). These issues are particularly pertinent to the representation of decision variables in the orbitofrontal cortex (OFC) – an area implicated in economic choices. Here we present a new algorithm to assess whether a neuronal representation is categorical or mixed, and to identify the encoded variables if the representation is indeed categorical. The algorithm is based on two clustering procedures, one variable-independent and the other variable-based. The two partitions are then compared through adjusted mutual information. The present algorithm overcomes limitations of previous approaches and is widely applicable. We tested the algorithm on synthetic data and then used it to examine neuronal data recorded in the primate OFC during economic decisions. Confirming previous assessments, we found the neuronal representation in OFC to be categorical in nature. We also found that neurons in this area encode the value of individual offers, the binary choice outcome and the chosen value. In other words, during economic choice, neurons in the primate OFC encode decision variables in a categorical way.

**Author Summary:** Mental functions such as sensory perception or decision making ultimately rely on the activity of neuronal populations in different brain regions. Much research in neuroscience is devoted to understanding how different groups of neurons support specific brain functions by representing behaviorally relevant variables. In this respect, one important question is whether individual neurons represent single variables, or linear combination of variables. Here we developed a new algorithm to assess this general issue. We then used the algorithm to examine neurons in the orbitofrontal cortex (OFC) recorded while non-human primates performed economic decisions. We found that different neurons represent different variables (categorical encoding). Specifically, neurons in the OFC encoded the value of individual offers, the binary choice outcome, and the chosen value. The present results support the hypothesis that economic decisions are formed within the OFC.

## Introduction

A recurrent question in systems neuroscience is that of understanding what variables are encoded by a given population of neurons. Addressing this issue is a prerequisite to understand what role neurons play in functions such as sensation or decision making. In a typical experiment, animal subjects perform some task, behavioral conditions vary along one or more dimensions, and the corresponding parameter(s) define variables potentially encoded by neurons in some brain area. In first approximation, if firing rates vary systematically with a variable, it can be said that neurons encode or represent that variable. Building on this concept, countless studies shed light on the neural substrates of sensory, associative and motor processes. Importantly, identifying the variables encoded by a given population can sometimes be challenging due to the trial-by-trial variability of neuronal firing rates combined with three other factors. First, different neurons, even in close proximity to one another, may encode different variables, and the number of variables encoded by a neuronal population is generally not known. This situation may arise in any brain area, but is most typical for prefrontal regions. Second, different candidate variables potentially encoded by the neuronal population may be substantially correlated with one another. Third, the encoding of different variables may be categorical (different neurons encoding different variables) or mixed (individual neurons encoding a linear or non-linear combination of variables). These two encoding schemes are referred to as pure selectivity and mixed selectivity, respectively [1]. Of course, the encoding scheme adopted by any particular population is not known a priori.

All these issues are particularly pertinent to the representation of decision variables in the orbitofrontal cortex (OFC) – an area implicated in economic (or value-based) decisions [2,3]. In recent years, numerous studies have shown compelling evidence for mixed selectivity in lateral prefrontal regions [4–8], suggesting that mixed selectivity is the hallmark of neural systems supporting complex cognitive functions [1,9]. At the same time, several studies argued for categorical encoding of decision variables in the OFC. Concurrent results in this sense came from studies of economic decisions in non-human primates [10,11] and from studies of decision confidence in rodents [12]. In contrast with these observations, a recent study argued for mixed encoding of decision variables in the primate OFC [13] (more on this below). Importantly, the categorical nature of this representation is a key assumption underlying current neuro-computational models of economic decisions [14–21]. Given the importance of this matter, we set up to revisit the question of categorical versus mixed selectivity in the OFC using a new and more powerful statistical approach. Our goal was to develop a set of procedures (or algorithm) with four objectives in mind. First, the algorithm should assess the categorical versus mixed nature of a neuronal representation without committing to any particular set of variables. Second, if the encoding was indeed categorical, the algorithm should facilitate a quantitative comparison of multiple candidate variables potentially represented by the neuronal population. Third, the algorithm should operate seamlessly in cases where different variables encoded in the neuronal population are correlated. Fourth, the algorithm should be amenable to general use, for any neuronal population and any behavioral task.

To achieve our stated objectives, we considered the high-dimensional space defined by all the behavioral conditions occurring in the task (referred to as “trial types”). We noted that each neuronal response corresponds to one point in this space. Furthermore, after normalization, each response corresponds to one point on the hyper-spherical surface of unitary radius. Cast in this terms, the problem of assessing pure versus mixed selectivity maps onto a clustering problem defined on a high-dimensional hyper-spherical surface, which we resolve using a spherical k-means approach [22]. In our algorithm, the categorical versus mixed nature of the representation is assessed before defining any behavioral variable. The spherical k-means returns a number of clusters and their locations in the space of possible responses (i.e., the hyper-spherical surface). Furthermore, any variable possibly encoded in the neuronal population (i.e., any quantity systematically varied across behavioral conditions) also corresponds to a point on the hyper-spherical surface. Casting a wide net, we can generate a large number of variables potentially encoded by the neuronal population and thus identify the subset that minimizes the total distance from the clusters. Importantly, these procedures are completely general and do not depend on the nature of the task, except for the definition of candidate variables potentially encoded by the neuronal population.

The Results are organized as follows. The first section describes the juice choice experiments conducted in monkeys, the neuronal data set collected in OFC, and previous analyses of these data. The second section introduces the new algorithm. The third section demonstrates how the criteria previously used to assess the categorical nature of the neuronal representation in OFC [10] can, in some cases, lead to erroneous conclusions. The fourth section describes the results obtained by testing the new algorithm on a set of synthetic data. In the following section, we describe the results obtained by analyzing the actual OFC data with the new procedures [23]. In a nutshell, the results corroborate previous findings [11]. In the Discussion, we compare the present algorithm to other approaches proposed in the literature. We also emphasize that procedures presented here provide a general and powerful method to analyze heterogeneous populations of neurons.

## Results

### Data set and previous analysis

In the experiments, two rhesus monkeys performed an economic choice task [11,23]. In each session, the animal chose between two juices offered in variable amounts. The preferred and non-preferred juices were labeled juice A and juice B, respectively. A “trial type” was defined by two offers and a choice (e.g., [1A:3B, B]), and each session typically included 5-20 trials per trial type. Our data set included 1008 neurons. Neuronal spiking activity was recorded and processed with standard techniques (see Methods). For the analysis of how firing rates depended on the task variables, we defined several time windows aligned with respect to different behavioral events. For each trial type and each time window, firing rates were averaged across trials. A “neuronal response” was defined as the activity of one cell in one time window as a function of the trial type.

Our previous analyses proceeded as follows [11,23]. First, each neuronal response was tested with an ANOVA (factor trial type). Responses that passed a statistical criterion (p<0.001) were considered task-related and analyzed further. Our data set included 2047 task-related responses. Second, we defined a large number of variables potentially encoded by this population. We performed a linear regression of each response on each variable, from which we obtained the regression slope and the R^2^. If the regression slope was significantly different from zero (p<0.05), the variable was said to “explain” the response. Third, two procedures – stepwise and best subset – were used to identify a small set of variables that best explained the neuronal population. In a first study [11], both procedures identified variables *offer value A*, *offer value B*, *chosen value* and *chosen juice*. This result was replicated several times, including in the data set examined here [23]. Finally, each neuronal response was assigned to the selected variable that provided the highest R^2^. Two additional analyses were conducted to address the issue of mixed versus pure selectivity. First, for each neuronal response it was assessed whether adding a second variable to the regression (through a bi-linear regression) would significantly improve the fit. This analysis found that this was the case for only a small fraction of responses [11]. A second analysis quantified for each neuronal response and for each pair of selected variables the difference in the corresponding R^2^ (ΔR^2^), and examined the distributions of ΔR^2^ across the neuronal population. In general, these distributions presented a significant dip close to zero, indicating that variables were encoded in a categorical way [10].

The approach for data analysis summarized above has the advantage that it allows to examine a large number of variables in parallel without biasing the conclusions, and that it withstands situations in which candidate variables are highly correlated with one another. At the same time, this approach presents two limitations. First, the analyses require to first define candidate variables, then identify the most explanatory ones, and finally assess whether the encoding is categorical. In contrast, it would be preferable to assess whether the encoding is categorical without committing to any particular variable or set of variables, and only later define variables that best capture each category of responses. Second, there are situations in which the argument for categorical encoding based on the distribution of ΔR^2^ is not valid (more on this below). The algorithm presented in this study addresses these limitations.

### Detection of categorical encoding using spherical clustering

To detect categorical encoding, we devised an algorithm that combines a clustering procedure partitioning neural responses based only on their spatial configuration with one that starts from a particular set of variables. In essence, the idea is to select a set of variables that best represents the spatial configuration of neural responses in the high-dimensional space of trial types.

**Fig 1** illustrates the algorithm for a 3-dimensional space (i.e., 3 trial types). Each data point represents a neuronal response (i.e., the activity of one cell, in one time window, averaged across trials for each trial type). Neuronal responses are first centered and normalized. This transformation places neuronal responses on a spherical surface of unitary radius. This data set undergoes two separate procedures for spherical clustering. First, the spherical k-means procedure, which does not assume any particular variable and yields a partition of the neural activity points based solely on the configuration of points in the high-dimensional space of trial types. For any number of clusters, this procedure alone reveals the categorical versus mixed nature of the neuronal data. Second, the variable-centroid clustering, which starts from a particular subset of variables (iteratively chosen from a large set of candidate variables; see **Table 1**). Notably, each variable corresponds to a point on the spherical surface. Thus, the subset of variables defines a corresponding number of cluster centroids, and we assign each neural response to the closest centroid. Each of these two clustering procedures (spherical k-means and variable centroid clustering) returns a partition of the population of neural responses. Importantly, the number of clusters is not known a priori. Furthermore, for any such number, there are many possible subsets of variables. We thus want to identify the subset of variables that best describes the neuronal data. As a measure of similarity between the two partitions, we use the adjusted mutual information [24]. Thus, we repeat the spherical k-means and the variable centroid clustering procedures for various number clusters and subset of variables. The variables that best match the non-committed spherical k-means partition are identified as encoded by the neuronal population.

**Table 1:**
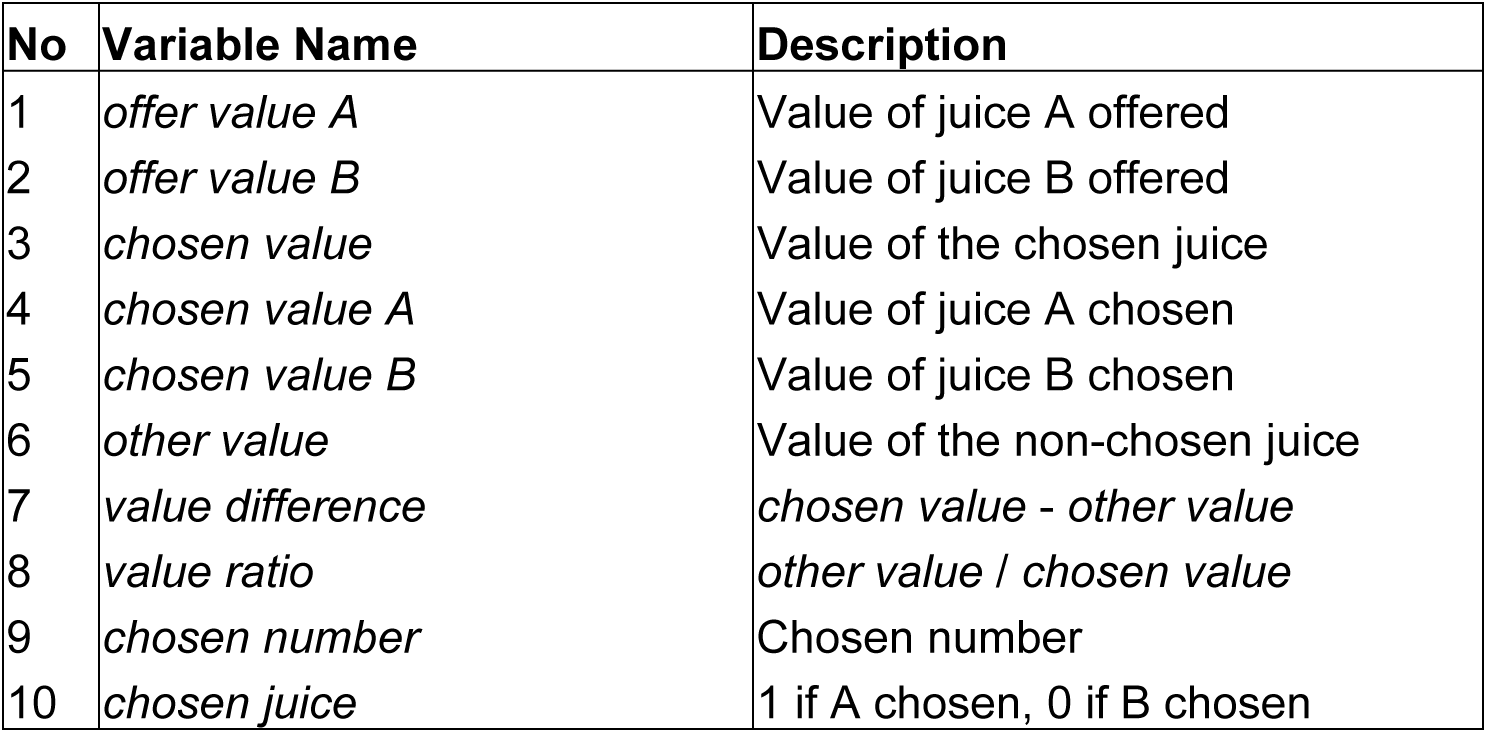
Considered list of variables that are potentially encoded by the population.

| No | Variable Name | Description |
| --- | --- | --- |
| 1 | *offer value A* | Value of juice A offered |
| 2 | *offer value B* | Value of juice B offered |
| 3 | *chosen value* | Value of the chosen juice |
| 4 | *chosen value A* | Value of juice A chosen |
| 5 | *chosen value B* | Value of juice B chosen |
| 6 | *other value* | Value of the non-chosen juice |
| 7 | *value difference* | *chosen value - other value* |
| 8 | *value ratio* | *other value / chosen value* |
| 9 | *chosen number* | Chosen number |
| 10 | *chosen juice* | 1 if A chosen, 0 if B chosen |

**Fig 1:**
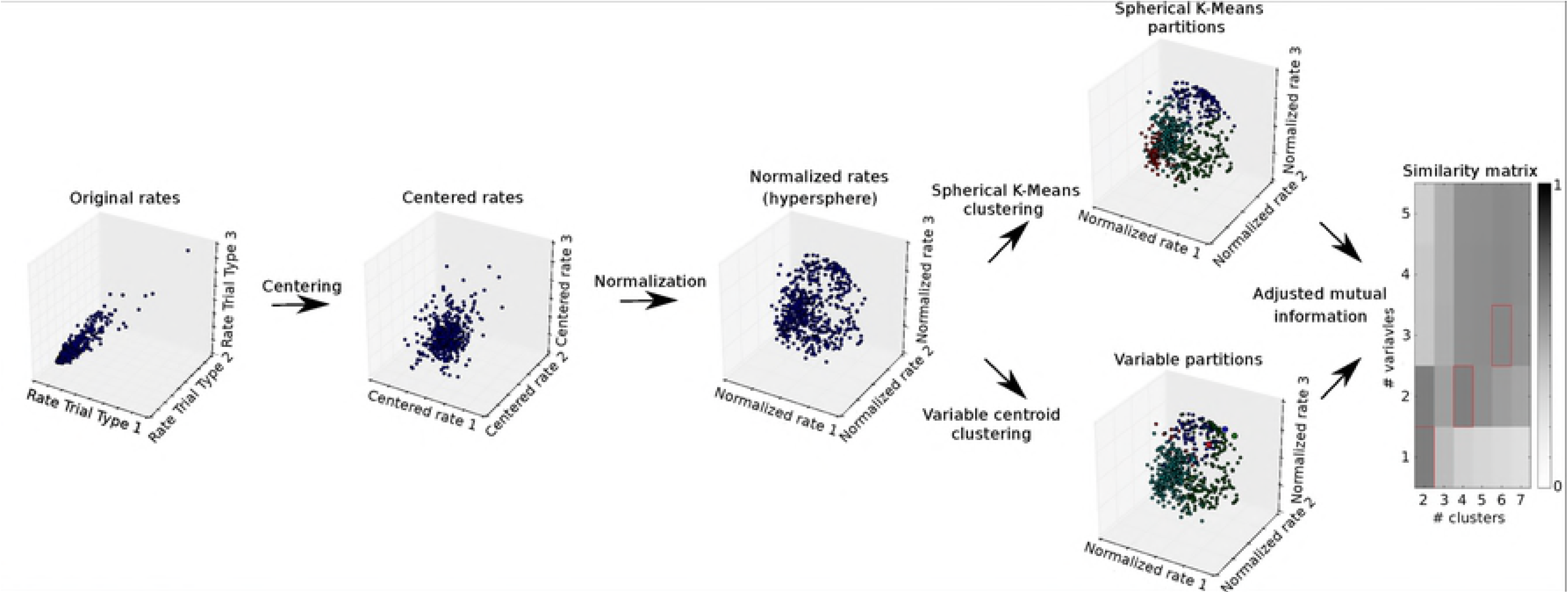
Illustration of the procedure for category discovery. For the original rates (left), each axis of the space denotes the firing rate of the cells in the various trial types. Each data point in this space represents a cell. For illustration, we present only 3 of the 9 dimensions. The original rates are then centered and normalized to unit length. The normalization effectively moves the points to the surface of a hyper-sphere. The points are then clustered using spherical k-means for a given number of clusters and centroid clustering for given variables. In the illustration, variables are represented as larger points. The resulting partitions are compared using the adjusted mutual information measure as a function of the number of clusters and number of variables (right).

### Limits of previous approaches

In previous work, the categorical nature of the neuronal representation in OFC was assessed through the analysis of the distribution of ΔR^2^ [10]. We will now demonstrate that this criterion can sometimes lead to erroneous conclusions. To do so, we construct two synthetic data sets and we show that the ΔR^2^ analysis fails while the spherical k-means reveals the true nature of the data. Again, each neural response is a point on the hyper-spherical surface of the a high-dimensional space defined by the trial types, and variables correspond to points on this surface.

To simulate different neuronal populations, we generated distributions of points on the hyper-spherical surface.

**Fig 2A** illustrates the first example. Here data points form two clusters: a circular cluster close to the spherical pole and a banana-like cluster along the equator. Importantly, the distribution used to generate the banana cluster was uniform on a banana domain (no intrinsic dip). We now examine the situation in which the analyst identified the wrong variables, shown as large circles in **Fig 2A**. We assume that the analyst correctly identified the pole variable, but erroneously selected two variables located at the opposite tips of the banana cluster. As illustrated in **Fig 2B**, the distribution of ΔR^2^ between the two banana variables has a significant dip around zero (Hartigan’s dip test, p<0.001) suggesting that the two variables are categorically distinct.

**Fig 2:**
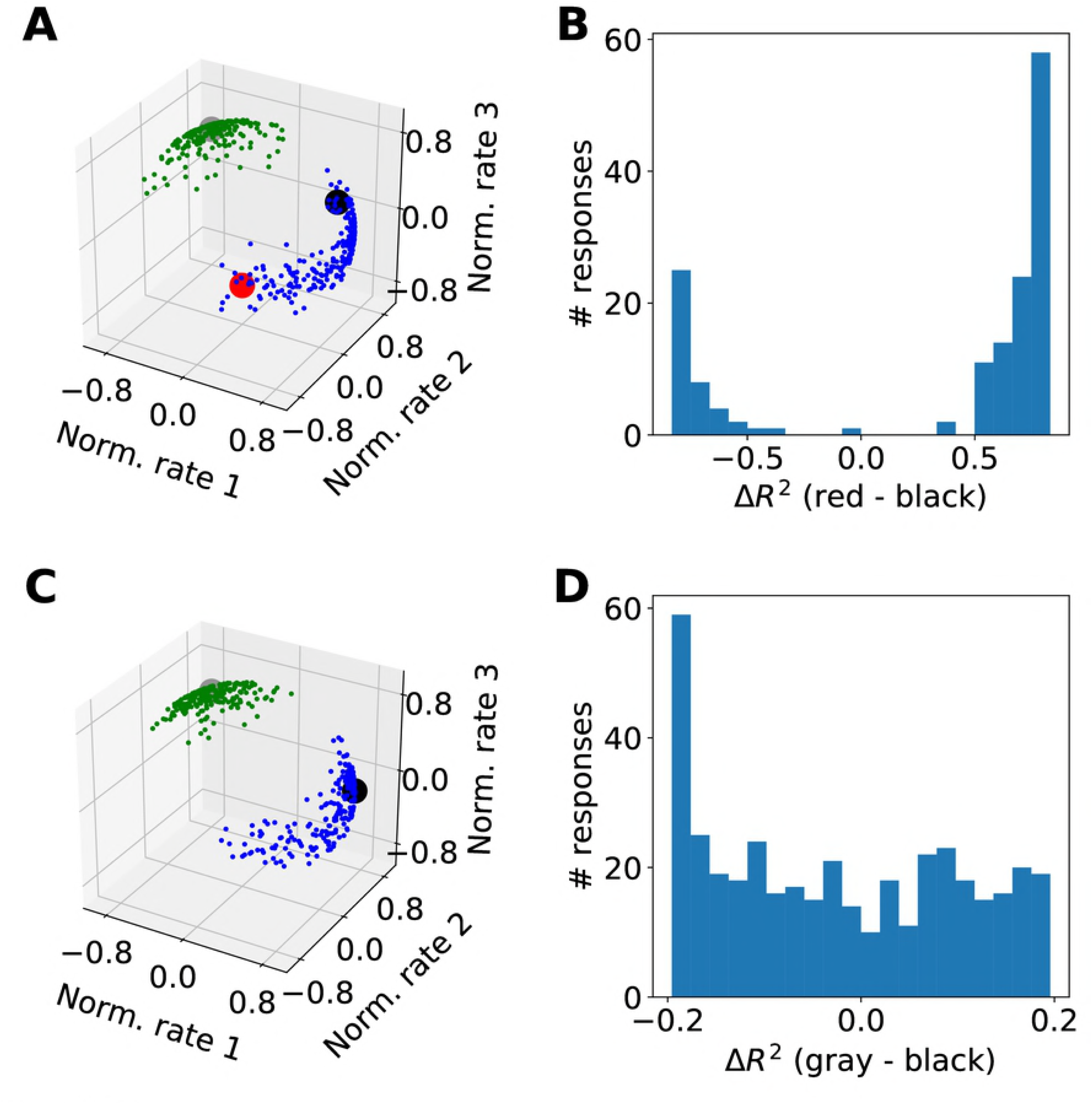
Two examples of how the ΔR^2^ metrics can fail. (A,B) A dip in the distribution of ΔR^2^ does not necessarily imply categorical encoding. The clustering algorithm yields two clusters. However, the analyst might have erroneously concluded that there are three variables, including two variables located in the tips of the banana cloud (red and black). The dip in the ΔR^2^ histogram suggests that these two variables are encoded by categorically distinct populations, but this is in fact not the case. (C,D) Categorical encoding does not always result in a dip in the distribution of ΔR^2^. In this case, we assume that the analyst correctly concluded that there are two variables, but might have defined these variables such that one is on the north pole (gray) and the other is on the east end of the banana cluster (black). Inspection of the ΔR^2^ histogram does not reveal any dip. The reason is that data points on the west end of the banana cluster are equally far from the two variables.

However, this suggestion is at odds with the ground truth. The dip in the distribution if ΔR^2^ is due to the presence of the third cluster, because some of the data points in the banana are closer to the pole variable than to either of the banana variables. Hence, a dip in the distribution of ΔR^2^ does not necessarily imply that the corresponding variables are encoded by categorically distinct groups of neurons. Importantly, the spherical k-means clustering correctly identifies the presence of two clusters (colors illustrate the k-means partitioning).

The second example makes the converse point, namely that a unimodal or uniform distribution of ΔR^2^ does not necessarily imply mixed selectivity. One obvious reason why this is the case is that absence of evidence is not evidence of absence; here we illustrate a more subtle issue. We consider the same clusters defined above. In this case, we assume that the analyst correctly identified two variables, one in the pole cluster and one in the banana cluster. However, we assume that the variable in the banana cluster is off center (**Fig 2C**). As illustrated in **Fig 2D**, the resulting ΔR^2^ histogram does not present a dip (Hartigan’s dip test, p=0.313), even though the two clusters are categorically separated. Importantly, the spherical k-means clustering correctly identifies the two clusters.

In conclusion, a dip in the distribution of ΔR^2^ is neither sufficient nor necessary to assess the categorical nature of a neuronal representation. In general, such assessment requires the examination of the spatial distribution of data points in a high-dimensional space, using an approach such as the spherical k-means clustering.

### Analysis of synthetic data

We considered several clustering procedures, and wanted to validate our algorithm to assess the categorical versus mixed nature of a neuronal representation on data for which we could control the ground truth. Thus, we generated synthetic populations of neural responses with and without specific categorical structure, and applied clustering algorithms to these synthetic data.

For the real data, the experiments included 9 or 10 trial types, resulting in 9- or 10-dimensional neuronal responses, represented as points on the unitary hyper-sphere in 9 or 10 dimensions. (see section *Data set and previous analysis*). To generate synthetic neuronal responses with categorical nature, we randomly generated 9-dimensional points on the hyper-spherical surface clustered in the vicinity of selected variables (see Methods). We then analyzed these synthetic data sets with a wide range of clustering algorithms, including centroid-based clustering methods (mini-batch k-means, spherical k-means), hierarchical clustering methods (Ward, agglomerative clustering, Birch), and a graph-based clustering method (spectral clustering) [25–30]. To estimate the performance of these algorithms, we used silhouette plots, which are a common method to assess the goodness of clustering partitions [31]. For each data point X (here X is a normalized neuronal response), the silhouette value quantifies the mean distance between X and other data points in the same cluster, and compares it to the mean distance between X and data points in the nearest other cluster. The greater the silhouette value, the better the clustering. A negative silhouette value indicates that X was assigned to the wrong cluster, since X is closer to the nearest other cluster.

**Fig 3** shows the silhouette plots obtained for the various clustering algorithms. We found that the hierarchical clustering methods (Ward, Agglomerative, Birch) produced the greatest number of negative silhouette values. Spectral clustering produced slightly less negative silhouette values than Ward as the best hierarchical clustering method. The centroid-based methods had no (spherical k-means) or very few (mini-batch k-means) negative silhouette values and many large silhouette values, suggesting that these methods found the most consistent clustering partitions. The silhouette analysis further suggests that the spherical k-means clustering is best suited for categorical data lying on a hyper-sphere. We also compared the silhouette plots on real data recorded from OFC. We found that spherical k-means had the smallest number of negative silhouette values, confirming the results from synthetic data. Of note, the superior performance of spherical k-means might be due to the fact that this algorithm explicitly considers the hyper-spherical structure of the data. Hence, we used spherical k-means for clustering in the remainder of this study.

**Fig 3:**
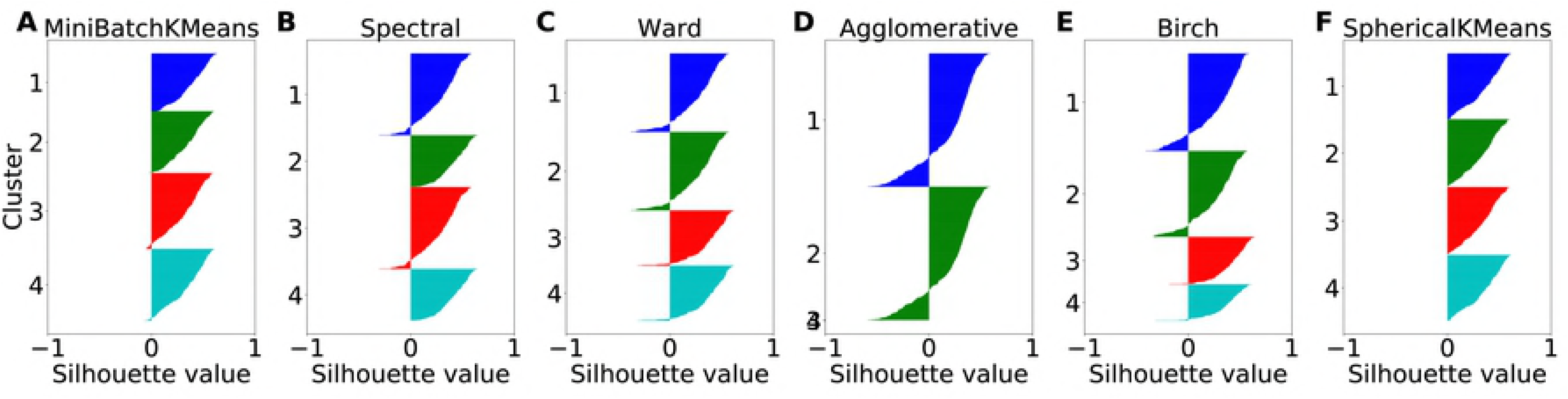
Silhouette comparison of clustering algorithms on synthetic categorical data. Synthetic data consist of firing rates from a total of 400 simulated cells representing the variables chosen value, offer value A, offer value B and chosen juice (100 cells each). Independent Gaussian noise with a standard deviation of 0.25 and a mean given by the variable rates was used to simulate the activity of a cell. Each color corresponds to one cluster. Clustering algorithms were Mini-Batch k-means (A), Spectral Clustering (B), Ward (C), Agglomerative Clustering (D), Birch (E) and Spherical k-means (F). The number of clusters was fixed to 4.

We next compared the spherical k-means silhouette plots for categorical synthetic data with those for non-categorical data (**Fig 4**). To simulate neural responses without specific categorical structure, we generated points uniformly on the hyper-spherical surface. We then varied the number of clusters between 2 and 7. We did not expect to find negative silhouette values for these data, because negative values indicate data point assignments to wrong clusters. Such mis-assignments cannot occur without any cluster structure in the data. Indeed, we did not find any negative silhouette values, neither for categorical data (**Fig 4 A-F**) nor for non-categorical data (**Fig 4 G-L**). However, while for categorical data the silhouette values in each cluster were dominated by large values yielding convex plots, the silhouette values for non-categorical data were dominated by small positive values yielding concave plots. Such concavity clearly indicates lack of cluster structure and allow to discriminate between categorical data and non-categorical data [31].

**Fig 4:**
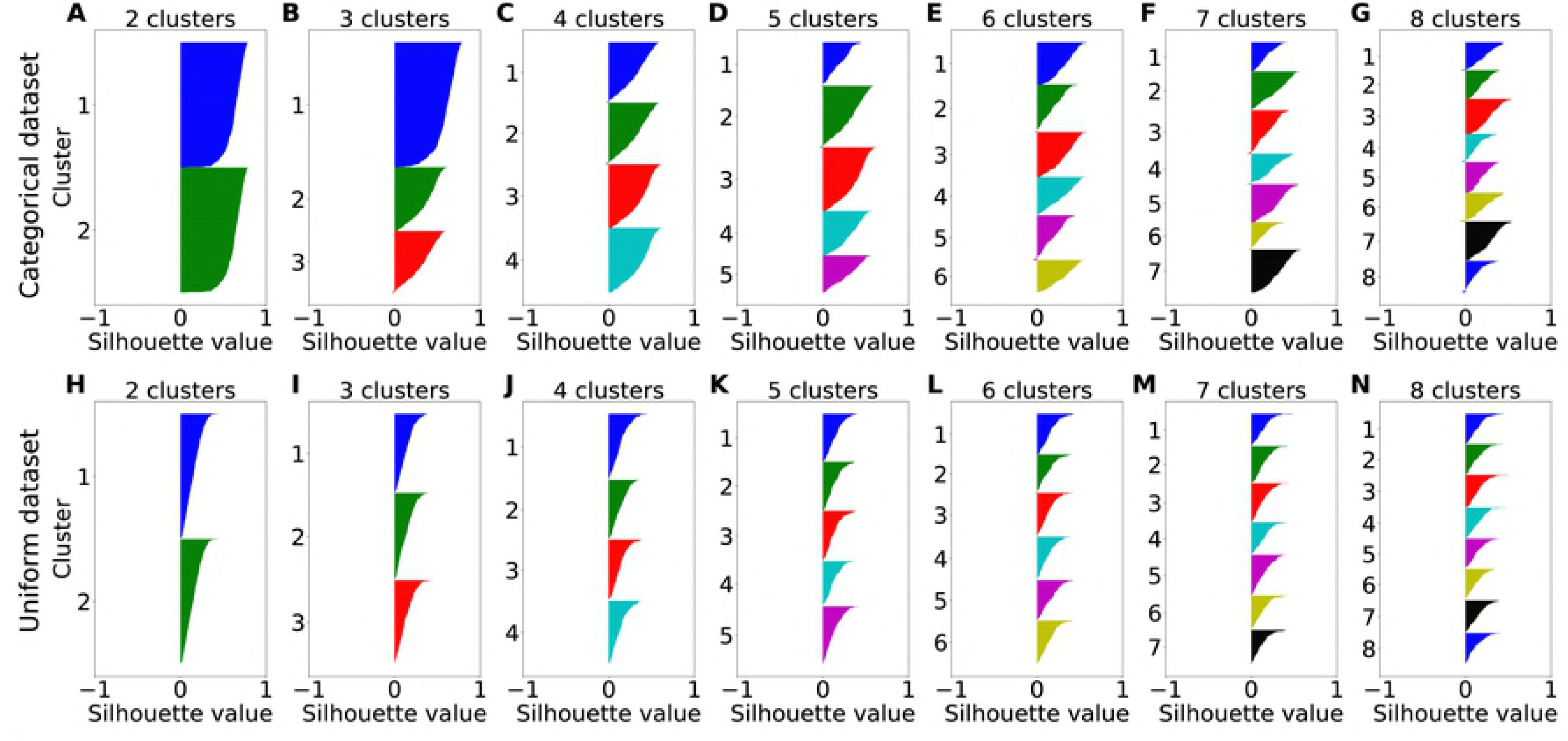
Silhouette comparison of spherical k-means clustering for different numbers of clusters on synthetic data. Synthetic data were either categorical (top row) simulated like in Figure 3 or non-categorical consisting of 400 samples uniformly distributed over the unit hyper-sphere (bottom row). For each data set, the number of clusters was varied between2 and 8 (A-G) and (H-N) respectively. Each color corresponds to one cluster.

While the silhouette analysis provides a simple way to evaluate the assignments of data points to clusters, it does not immediately associate particular variables with clusters. To establish this relation, we devised a comparative clustering method. In addition to spherical k-means, we performed a centroid-based clustering where the centroids were defined by a particular set of variables. We refer to this procedure as “variable-centroid clustering”. We assigned each data point to the nearest centroid on the sphere (see Methods). We then compared the clusters obtained from spherical k-means to the clusters obtained from variable-centroid clustering, and quantified their similarity for different sets of variables.

Quantifying the similarity of two clustering partitions is non-trivial because similarity should be invariant for cluster relabeling. Many measures of similarity have been proposed [24,32,33]. Here we tested three measures of similarity based on mutual information, which are founded on information theory and naturally satisfy our desiderata. Specifically, we tested mutual information (MI), normalized mutual information (NMI) and adjusted mutual information (AMI).

MI quantifies the information one clustering partition provides about another clustering partition; NMI normalizes MI yielding values between 0 and 1; AMI additionally corrects for the agreement expected by chance. We compared the performance of these candidate measures of similarity using our synthetic categorical and non-categorical data sets. **Fig 5** shows the results obtained for each measure as a function of the number of clusters specified in the spherical k-means algorithm and the number of variables defined in the variable-centroid clustering. For each number *n* = 1, 2, … of variables, we tested all of the possible sets of *n* variables, and we identified the set providing the maximum similarity. We used exhaustive search for this purpose (see Methods).

**Fig 5:**
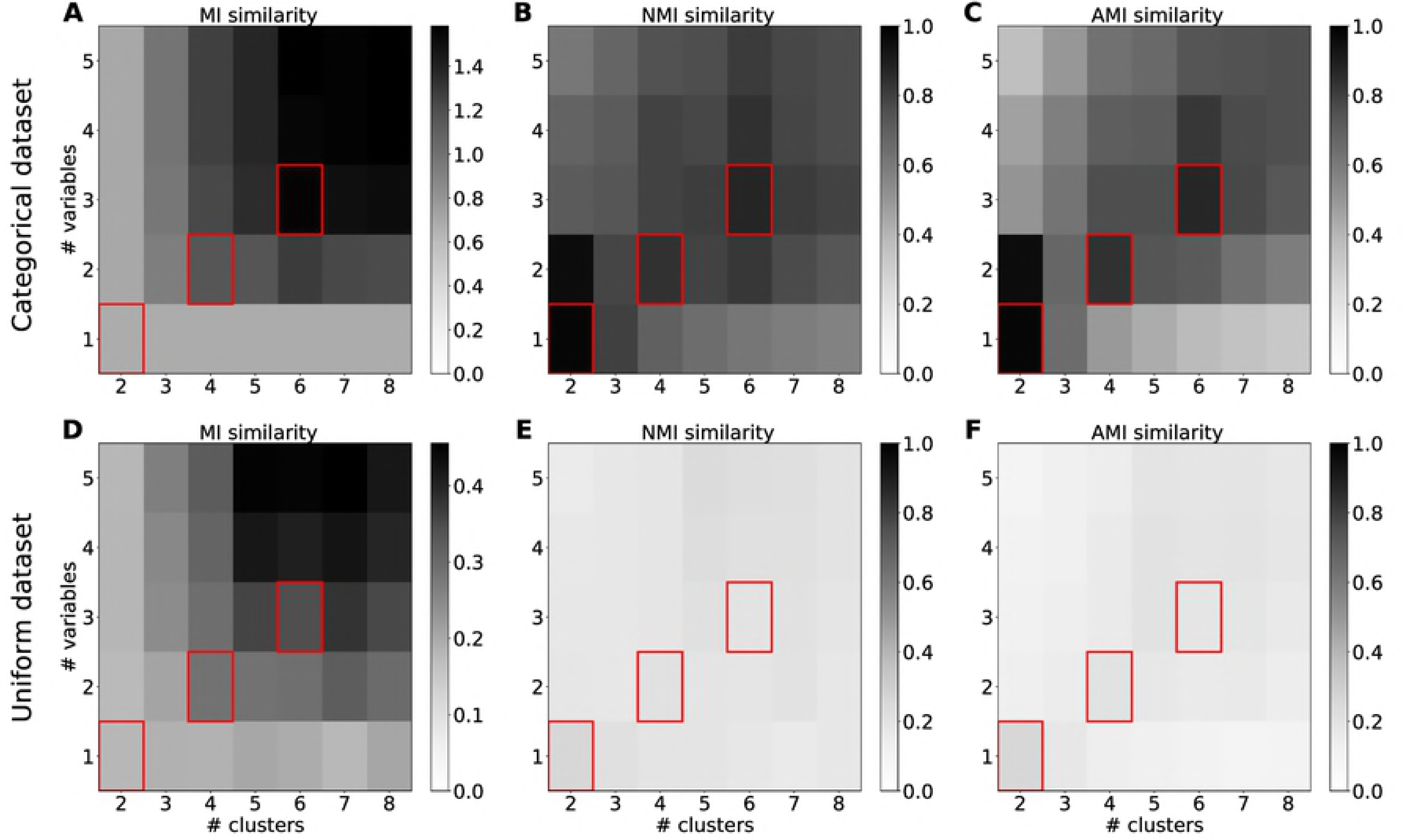
Comparison of different cluster similarity measures for spherical k-means partitions on synthetic data. Data were either categorical (top row) or non-categorical (bottom row) and simulated like in Figures 3 and 4. The similarity measure was either mutual information (A, D), normalized mutual information (B, E) or adjusted mutual information (C, F). The gray scale indicates the strength of similarity for given number of clusters and number of variables. Corresponding numbers of clusters and numbers of variables are marked in red.

For both categorical and non-categorical data, MI tended to increase with the number of clusters and variables (**Fig 5A, 5D**). This was expected since additional clusters and variables can convey more information about each other. Importantly, MI increased to ~0.5 bits even for non-categorical data, highlighting the necessity for normalization. The additional normalization in NMI yielded clear peaks for categorical data and mostly flat values for non-categorical data.

This made it easy to discriminate between categorical and non-categorical data based on NMI. Additionally, the peaks indicated corresponding numbers of clusters and variables where n variables correspond to 2n clusters. This was because the reflection of data points on the hyper-sphere (see Methods) produced twice the number of clusters. This reflection also facilitated the separation of the data points into two clusters for both one and two variables. For this reason, the very strong peaks for two clusters should be ignored. The results obtained for AMI (**Fig 5C, 5F**) were very similar to those for NMI. The peaks for corresponding cluster numbers and variable numbers were slightly sharper for AMI. For this reason, we selected AMI as our similarity measure for the analysis of real neural data recorded from OFC.

In conclusion, the analysis of synthetic data with known ground truth showed that a combination of spherical k-means clustering and variable-centroid clustering compared with AMI provided the most powerful approach to assess the categorical nature of neuronal representations and to identify the encoded variables.

### Analysis of neuronal data

We analyzed neuronal activity recorded from OFC during experiments in which monkeys chose between different juice types (see section *Data set and previous analysis*). In total, we analyzed 9 neuronal pools, each including 139-536 neuronal responses (see Methods). Applying to each pool the same comparative clustering procedure with spherical k-means and AMI used for synthetic data, we obtained silhouette plots and a similarity profile for the neuronal data.

We varied the number of clusters between 2 and 7 and found clusters with convex silhouette plots indicating categorical data (**Fig 6A-F**). Moreover, the almost complete absence of negative silhouette values indicated that the spherical k-means found consistent partitions for different number of clusters. The normalized neuronal data contains 9-10 dimensions (corresponding to trial types) which are hard to visualize. In **Fig 7** we illustrate the 2-dimensional projections of a data set consisting of 9-dimensional response. Four clusters are color-coded. Even though the clusters in this representation are partly overlapping, there is a clearly discernible structure. For a qualitative assessment of the results, we examined the response prototypes defined by the centers of individual clusters. In general, the response prototypes obtained for n = 3, 4, 5 closely resembled the neuronal responses illustrated in previous studies [11,23]. One example is illustrated in **Fig 8**. In other words, the clusters obtained from the spherical k-means qualitatively validated previous conclusions.

**Fig 6:**
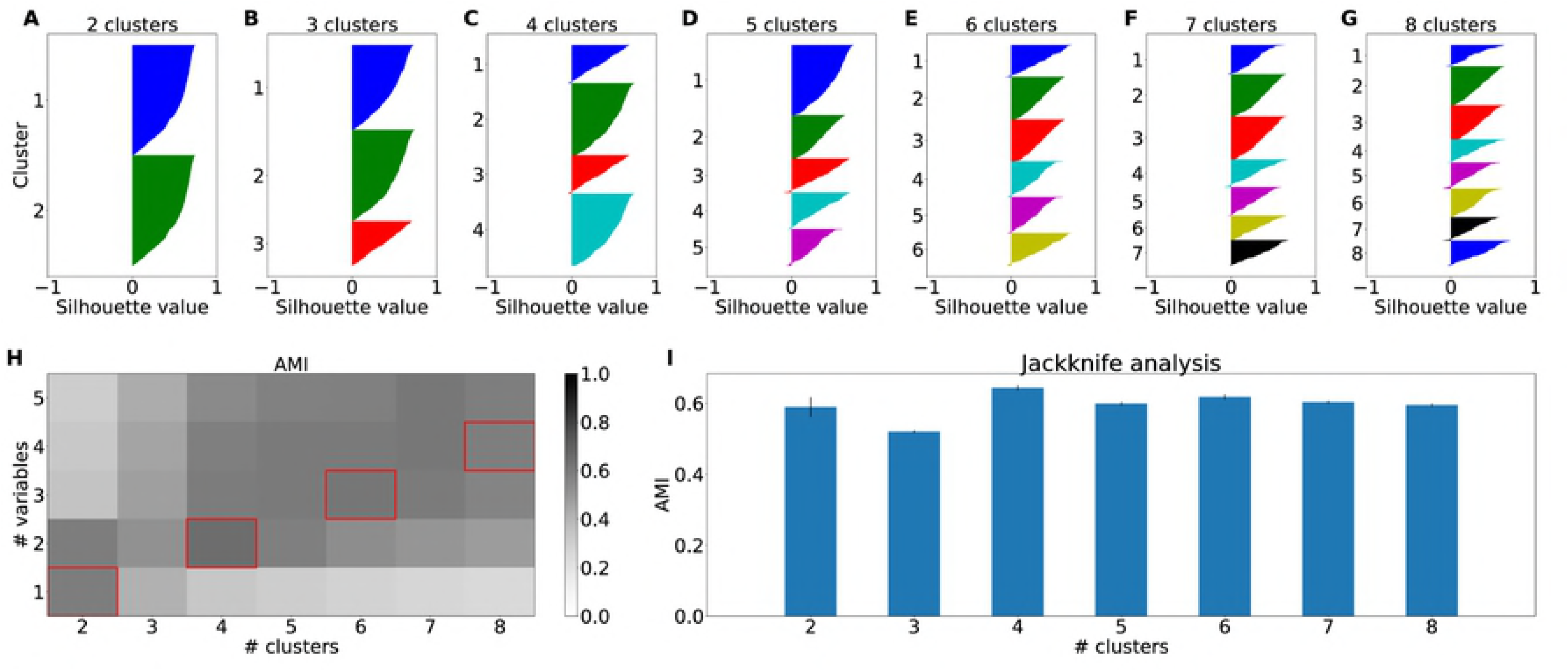
Cluster results for real data recorded from macaque orbitofrontal cortex. (A)-(F) Silhouette plots for the spherical k-means partitions of one example pool. Each color corresponds to one cluster. The number of clusters was varied between 2 (A) and 8 (G). (H) Adjusted mutual information cluster similarity between spherical k-means clustering and variable-based centroid clustering as a function of the number of clusters and number of variables over all pools. Corresponding numbers of clusters and numbers of variables are marked in red. (I) Maximum adjusted mutual information for each number of clusters with Jackknife estimated standard errors.

**Fig 7:**
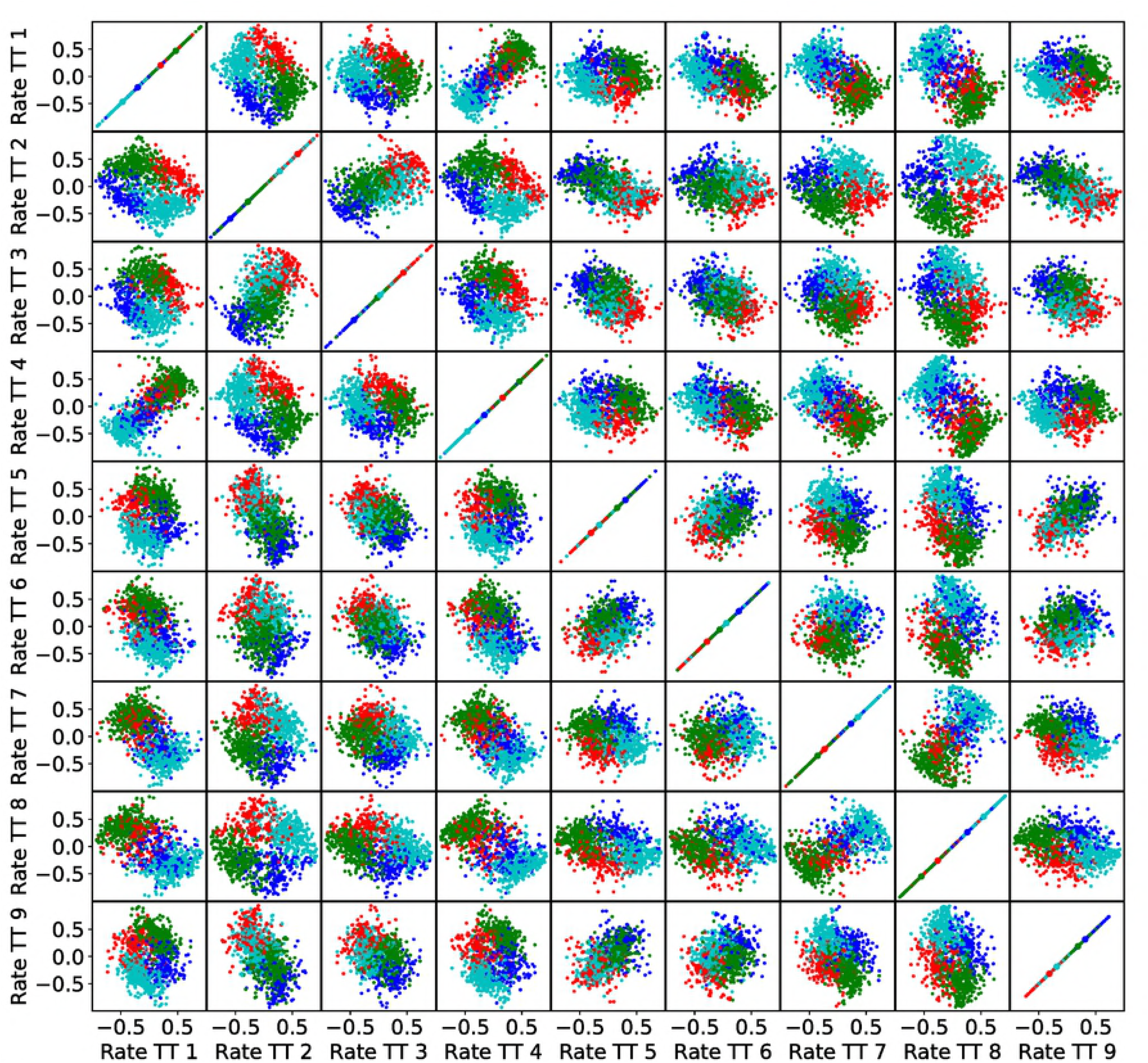
Visualization of four clusters in the 9-dimensional trial type space. Each color corresponds to one cluster. Each panel shows the centered and normalized firing rates of a pair of trial types and each point in a panel represents a cell. Cluster centers are marked with black circles.

**Fig 8:**
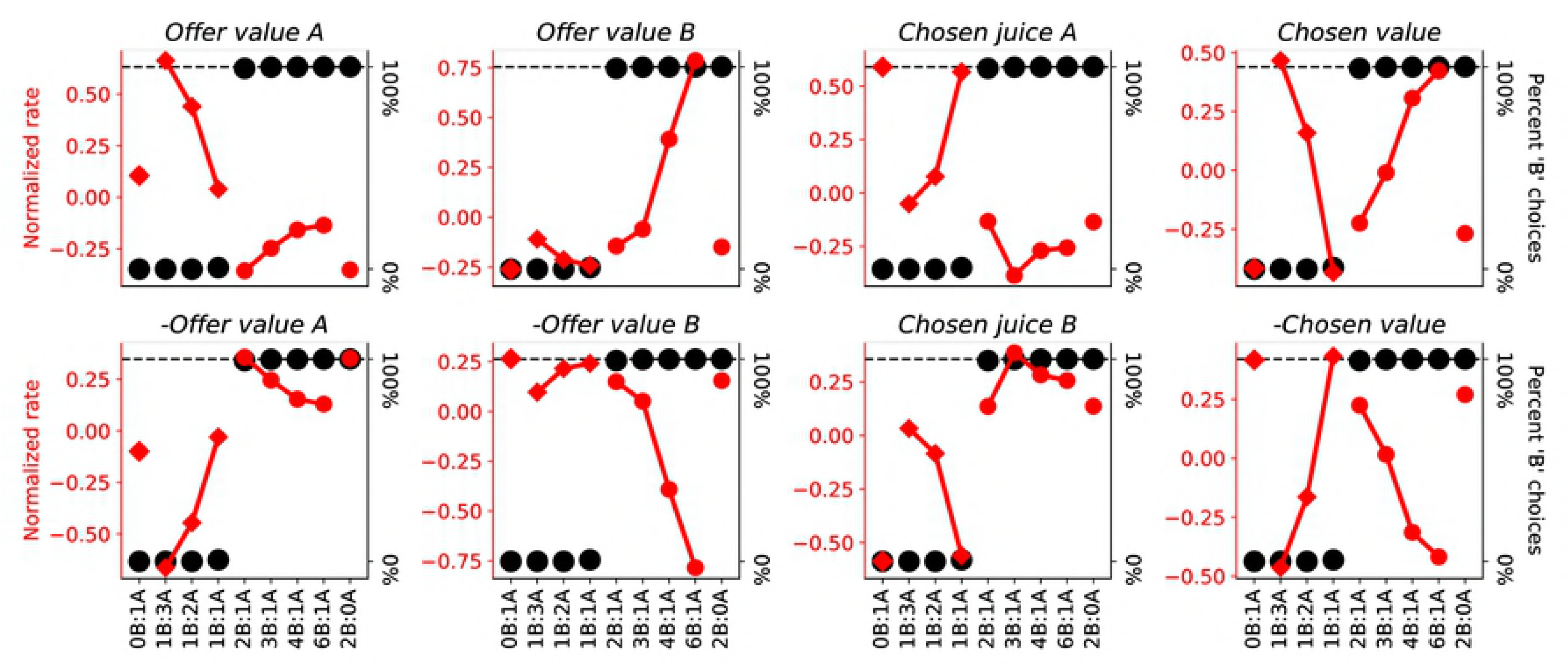
Tuning curves of response prototypes as defined by 8 cluster centers. The x-axis represents offer types ranked by the ratio #B:#A. The y-axis in red represents normalized response rates. The y-axis in black shows monkey behavior. Encoded variables are denoted in the panel titles. Red diamonds represent the responses to chosen juice A whereas red dots represent the responses to chosen juice B. The separate red diamond and red dot show forced choices.

For a quantitative assessment, we used AMI. Comparing the k-means clusters and the variable-centroid clusters, we found similarity peaks for particular combinations of cluster and variable numbers (**Fig 6G**). These peaks resembled those obtained for synthetic data, providing further evidence for the categorical structure of the neural data. To analyze in more detail the clusters and variables yielding maximum AMI we performed a Jackknife analysis (see Methods). This procedure allowed us to estimate the standard error of AMI values for a given number of clusters (**Fig 6H**). Excluding the peaks for 2 clusters, we obtained the highest AMI values for 4 clusters and 2 variables. The AMI for this combination of variables and clusters was significantly greater than the second largest AMI (Wilcoxon rank sum test, p<0.001).

**Table 2** summarizes the results of our analyses. For 4 clusters and 2 variables, the algorithm selected variables *chosen value A* and *chosen value B* for all neuronal pools. For 6 clusters and 3 variables, variables *offer value A*, *offer value B* and *chosen juice* were selected for all pools.

**Table 2:** Selected variables.

| #variables | 1 | 2 | 3 | 4 | 5 |
| --- | --- | --- | --- | --- | --- |
| **Selected variables** | *chosen value* | *chosen value A*<br>*chosen value B* | *offer value A*<br>*offer value B*<br>*chosen juice* | *offer value A*<br>*offer value B*<br>*chosen juice*<br>*chosen value* | *offer value A*<br>*offer value B*<br>*chosen juice*<br>*chosen value*<br>*value ratio* |

For 8 clusters and 4 variables, the algorithm selected variables *offer value A*, *offer value B*, *chosen value* and *chosen juice*. Note that these are the same variables identified in previous studies [11,23]. For 10 clusters and 5 variables, the algorithm selected these same variables plus the variable *value ratio* (= other/chosen value). The substantial consistency in the variables identified with increasing numbers of clusters indicates that the results are very robust.

## Discussion

We presented a new algorithm to assess whether a neuronal representation is categorical or mixed, and to identify the encoded variables if the representation is indeed categorical. The method involves two steps. First, we cluster the data without committing to any particular variable. Second, we match clusters with a set of candidate variables. Quantifying similarity between the clusters of the two steps makes it possible to identify the variables most consistent with the neuronal data. This new method overcomes limitations of previous approaches, and is widely applicable. In this study, we tested the algorithm on synthetic data and on neuronal data recorded in the primate OFC during economic decisions. With respect to the latter, the most notable result is that we found the neuronal representation in OFC to be categorical in nature. This result confirms previous assessments of this same data set [11,23], and the results obtained by other research groups [12]. We suggest that the categorical nature of the neuronal representation sets apart OFC from other prefrontal regions, where task-relevant variables are encoded in mixed representations [4–8].

In addition, our algorithm identified a set of variables encoded in OFC. The variables most reliably detected – *offer value A*, *offer value B*, *chosen value* and *chosen juice* – coincide with those identified in previous studies [10,11]. One difference concerns the number of variables. Previous work identified 4 variables imposing a criterion on the marginal explanatory power (i.e., each additional variable should explain ≥5% responses) [11,23,34]. In contrast, the AMI criterion establishes the optimal number of variables as 2. Several elements may explain this finding.

The AMI procedure penalizes the addition of further variables and thus tends to provide a conservatively small number of clusters. Concurrently, the variables encoded in OFC are substantially correlated in the experiments [11]. Geometrically, this means that the centers of different clusters are close to each other on the hyper-sphere, and not distributed randomly as might implicitly be assumed. Exacerbating this issue, in our data, neuronal responses encoding the *chosen value* have some additional jitter, because the relative value of two juices varied to some extent from session to session. This fact effectively broadened the corresponding cluster on the hyper-sphere.

### Comparison with other approaches

In previous work, we assessed the categorical nature of the representation in OFC based on linear regressions and the analysis of the resulting R^2^ [10,11]. As discussed above, that approach has some limitations, addressed by the algorithm presented here.

Another clustering-based method to assess categorical encoding was recently proposed by Hirokawa et al [12]. Their data set was recorded from the rat OFC and included 42 conditions. Applying principal component analysis as a pre-processing step, they first reduced this data set to 21 dimensions. Using spectral clustering, they identified 9 clusters (the number of clusters was determined based on bootstrap stability). While there are clear similarities between their approach and ours, there are also notable differences. Both approaches are founded on clustering of pre-processed neuronal activity. Hirokawa and colleagues applied spectral clustering, while we applied spherical k-means. On simulated data, we compared silhouette plots of several clustering procedures and we found that spherical k-means performed best. Most importantly, our approach associates easily interpretable variables with the identified clusters by making use of two comparative clustering steps – spherical k-means and variable-centroid clustering. Spherical k-means operates without prior assumptions on particular variables while variable-centroid clustering can be thought of as a cluster representation of a set of variables. By selecting the set of variables most similar to the assumption-free clusters, we obtain unbiased representations of neuronal categories.

Interestingly, both our results and the results of Hirokawa et al [12] are in contrast with those of a study by Blanchard et al, who argued that the neuronal representation in OFC is mixed and not categorical [13]. A close examination of this discrepancy highlights the advantage of assessing the categorical versus mixed nature of a neuronal representation without committing to any particular set of variables. Blanchard et al examined data from an experiment in which monkeys chose between two gambles. Apart from the stakes, which varied randomly from trial to trial, the two gambles differed qualitatively – one gamble was “informative”, meaning that the outcome (win or loose) would be revealed to the animal shortly after the choice and before the trial end; the other was “uninformative”, meaning that the animal would learn the outcome only at the end of the trial. Informative and uninformative gambles were associated with different colors, and the gamble informativeness consistently affected choices [35]. In their analysis, Blanchard et al [13] regressed each neuronal response recorded in OFC separately on the stakes and on the informativeness. They then examined the distribution of regression coefficients in 2D. Since the distribution was not condensed along the two axes, they concluded that there is no categorical tuning in OFC. The problem with this approach is that the neuronal representation may indeed be categorical, but the frame of reference and/or the variables encoded by the population may not be those tested in the analysis. For example, consider the hypothesis that in Blanchard's experiment there were four separate groups of cells encoding the value of the informative offer and the value of the non-informative offer, each with positive or negative sign. Such representation is categorical, not mixed. Yet, simple considerations show that Blanchard's analysis based on separate regressions on stakes and informativeness would fail to reveal the categorical nature of this representation, leading to the mistaken conclusion that the representation is mixed. To visualize this point, consider the clustering problem discussed in the present study. Choosing two specific variables is equivalent to choosing a particular plane and to projecting all the data set from the hyper-sphere on that plane. Unless the plane is defined in a sensible way, separate clusters will overlap and appear non-separable on the plane. In conclusion, it is preferable to assess the categorical or mixed nature of the representation without committing to a particular set of variables. As a second best, it is crucial to define variables in a sensible way, and to recognize that the conclusions drawn from regression analyses critically depend on the variables tested.

## Methods

### Experimental design and data set

The experimental procedures for data collection and preliminary data analyses have been described before [23]. Briefly, two monkeys participated in the study. All experimental procedures conformed to the NIH *Guide for the Care and Use of Laboratory Animals* and were approved by the Institutional Animal Care and Use Committee (IACUC) at Washington University in St Louis (protocol #20140031). Throughout the study, the animal health was overseen by a veterinary staff. Before training, a head restraining device and a recording chamber were implanted under general anesthesia (Isoflurane). Steps taken to increase the animal welfare included pair housing, cage enrichment, and usage of exclusively positive reinforcers.

In each session, a monkey chose between two juices (labeled A and B, with A preferred to B) offered in variable amounts. Each trial started with the animal fixating the center of a computer monitor. After 0.5 s, two sets of colored squares representing the two offers appeared on the two sides of the fixation point. For each offer, the color represented the juice type and the number of squares represented the juice amount. The animal maintained central fixation for a randomly variable delay (1–2 s), after which the fixation point was extinguished and two saccade targets appeared by the offers (go signal). The animal indicated its choice with a saccade and maintained peripheral fixation for 0.75 s before juice delivery.

In this experiment, the same neuron was recorded during two subsequent blocks of trials. Juices offered in the two blocks could be the same or different [23]. For the purpose of the present analysis, we considered data in each trial block independently. Thus each neuron appears in the analysis twice and the term “session” refers to a block of trials. In each session, offered quantities varied from trial to trial. An “offer type” was defined by two offers (e.g., [1A:3B]). Different offer types were pseudo-randomly interleaved. Their frequency varied, but each offer type was typically presented at least 20 times in each session. A “trial type” was defined by an offer and a choice (e.g., [1A:3B,A]).

In each session, choices were analyzed with a logistic regression:

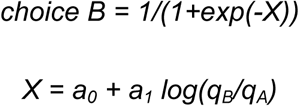

where *q_A_* and *q_B_* were the quantities of juices A and B offered to the animal. The relative value of the juices was inferred from the flex of the sigmoid and defined as *ρ = exp(-a_0_/a_1_)*.

Neuronal data were recorded from central OFC using standard techniques [23]. The analysis of firing rates was based on four primary time windows: post-offer (0.5 s after the offer), late delay (0.5–1 s after the offer), pre-juice (0.5 s before juice delivery), and post-juice (0.5 s after juice delivery). For each trial type and each time window, firing rates were averaged across trials. A “neuronal response” was defined as the activity of one cell in one time window as a function of the trial type. Task-related responses were identified with an ANOVA (factor trial type, p<0.001).

In preliminary work, we submitted the present data set to standard analyses for variable selection. In these analyses, we defined a large number of variables (**Table 1**), regressed each response on each variable, and used methods for variable selection to identify a subset of variables that best explained the population (see Results and [11]). These procedures replicated previous results, as neuronal responses were found to encode variables *offer value A*, *offer value B*, *chosen value* and *chosen juice* [11].

### Neuronal pools

The hyper-spherical clustering procedures introduced in this study require that different neuronal responses be defined on the same trial types (i.e., in the same space). Importantly, the offer types presented to the animal in our experiments could vary from session to session, although the same few sets of offer types were used repeatedly in many sessions. As a result, the entire data set could be divided in six groups of neuronal responses defined on the same trial types.

The variables included in the analysis are defined in **Table 1**. Of note, some variables (e.g., *chosen value*) were defined based on the relative value of the juices, which depends on the animal choices and thus varies somewhat from session to session. Ideally, the analyses described in this study would be conducted on pools of neuronal responses recorded in the same session, such that variables would be defined equally for all the responses. In contrast, our neurons were recorded in different sessions. Hence, we grouped responses in pools of similar relative values. For each group of neuronal responses recorded with the same trial types we examined the distribution of relative values. For five of the six groups, the distribution was bimodal. Hence, we split each of them in two and we removed outliers based on the inter-quartile range (IQR). In conclusion, our data set included 9 pools of neuronal responses recorded with the same trial types and similar relative values. (The remaining variability in relative values was effectively a noise factor that, if anything, made it more difficult to show categorical encoding.) Neuronal pools included 139-536 responses, and each pool was analyzed separately. When combining similarity values obtained for different pools, we weighted the similarities according to the number of neurons in the pool.

### Spherical representation of neuronal responses and variables

We represented neuronal responses as points in a high-dimensional space where each axis corresponds to a trial type. Raw neuronal responses were centered (by subtracting the mean firing rate across trial types) and normalized (imposing a unitary vector length). As a result, the neuronal population was constrained to the hyper-spherical surface of unitary radius. Similarly, for each variable we calculated a vector with elements given by the variable value in each trial type. We then centered and normalized the vector. Hence, each variable was represented as a point on the unitary hyper-spherical surface.

Previous work indicated that neuronal responses can encode a variable with positive or negative slope [11]. Hence, the sign of the normalized vector is ambiguous. For this reason, before conducting the clustering procedures, we mirrored each data point on the hyper-spherical surface. Resulting cluster centers were most of the time, but not always, symmetric when adding the mirror points. In principle, non-symmetrical clusters may be understood considering even and odd numbers of clusters. Symmetry implies an even number of true clusters, which is not necessarily the case. For instance, consider the case of the 3D sphere. If the raw data present only one cluster along the equator, adding mirror points will not generate a second cluster. If we run the algorithm imposing two clusters, the algorithm will place two cluster centers somewhere on the equator, but not necessarily on opposite ends. Now consider a situation where the raw data present a cluster along half of the equator and another cluster at one pole. Adding mirror points will result in one cluster along the equator and one cluster at each pole (3 clusters total). These examples demonstrate that mirroring does not necessarily induce an even number of clusters or symmetric cluster centers.

## Variable selection procedure

We selected variables by evaluating cluster similarity of partitions induced by a set of variables and of partitions obtained from spherical k-means clustering. The general algorithm for selecting the most informative set of variables works as follows:

- For given number of clusters, partition cells using spherical *k-means* clustering yielding partition *U* (see below)
- For given number of variables *n*, select variables by:
  ◦Iterate over different combinations *c* of *n* variables:
    ▪For given set of variables *c*, use each variable as a cluster center and cluster cells by means of proximity clustering yielding partition *V* (see below)
    ▪Evaluate similarity between partitions *U* and *V* using adjusted mutual information (AMI)
  ◦Select variable combination *c* that maximizes similarity

We clustered cells using the spherical *k-means* algorithm [26]:

1. Start with a partitioning 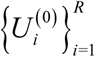 and the centroids 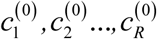 associated with the partitioning. Set the index of iteration *t* = 0.
2. For each normalized rate vector *x* find the centroids *c_i*(x)_* closest in cosine similarity to *x*, i.e.:

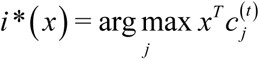 Next, compute the new partitioning 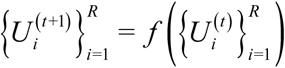 induced by the old centroids 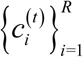:

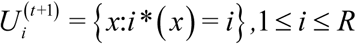
3. Compute new centroids corresponding to the partitioning computed for 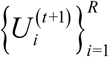:

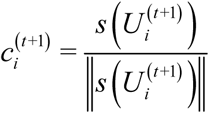

where 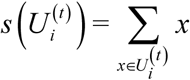
4. If 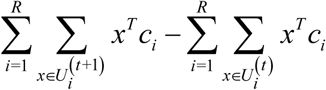 is greater than the tolerance 1e-4 (default value), increment *t* by 1 and go to step 2. Otherwise, stop.

For given variables, we partitioned cells using proximity clustering: for each cell, we calculated the cosine distance to each variable and assigned the cell to the variable with the smallest distance. Variables therefore became centroids of the clusters.

Our similarity measure between partitions is the adjusted mutual information:

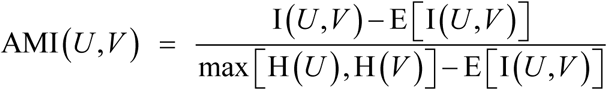

where *U* = {*U*_1_,…,*U_R_*} and *V =* {*V*_1_,…,*V_C_* denote two partitions of the same data, E denotes expectation of the mutual information over random partitions subject to having a fixed number of clusters and points in each cluster, H denotes entropy:

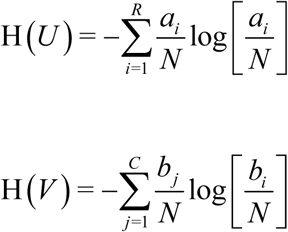

and I(*U*,*V*) denotes mutual information [36] between *U* and *V*:

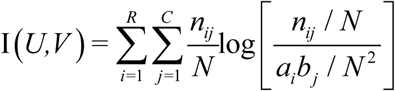

Mutual information was used because of its several advantages as metric for computing statistical associations between neural variables or between neural and behavioral variables, namely its ability to capture all forms of associations between such variables, including both linear and non-linear ones at all orders [37]. In the above equation for I(U;V), *n_ij_* denotes the number of objects that are common to clusters *U_i_* and *V_j_* and 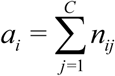 and 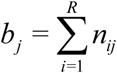.

Subtraction of the expectation values in the numerator and denominator adjusts the measure for chance and effectively corrects the positive bias of the measure. These terms can be calculated analytically [24]. In particular, we used the Python implementation *sklearn.metrics.adjusted_mutual_info_score* of the Scikit-learn package to calculate the AMI.

We checked all possible variable combinations (stopping at 5 variables) and collapsed the variables *offer value A* and *offer value B* to *offer value A|B* as well as *chosen value A* and *chosen value B* to *chosen value A|B* by pruning variable combinations that contained one but not the other of the collapsed variables. We then selected the variables and clusters with the greatest adjusted mutual information similarity.

## Jackknife estimates of standard error

We estimated standard errors of adjusted mutual information values by apply the Jackknife procedure over pools [38]. For a given number of clusters and a given jackknife subsample, we took the maximum AMI over the different numbers of variables. This yielded a (#clusters x #subsamples) matrix. We then collapsed the subsample dimension in two ways: 1) For a given number of clusters, we averaged over subsamples to get the mean AMI. 2) For a given number of clusters, we used the jackknife equation for standard deviation [38] to get an estimate of the standard error:

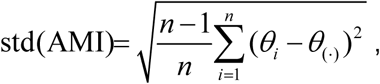

where *θ_i_* denotes the *i*-th AMI estimate and

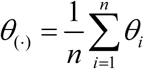

denotes the mean AMI.

## Generation of synthetic data

We generated two synthetic data sets to test our variable selection procedure: one with categories and the other without categories.

To generate the data set with categories, we selected four variables: *total value*, *offer value A*, *offer value B* and *chosen juice*. We represented each of these variables in the trial type space on the hyper-sphere as a 9-dimensional vector with unit length (see Section “Representation of cells and variables”). Then, for each of these variables, we generated 100 synthetic cell responses by adding independent Gaussian noise to each of the vector elements (zero-mean, standard deviation 0.25). Using this procedure, we obtained point clouds around each variable consisting of 100 points each. Finally, we moved the points to the unit hyper-sphere by normalizing the vector length of each point. Each point then represented the centered and normalized firing rates of a synthetic cell.

To generate a data set without categories, we drew 400 samples uniformly on the 9-dimensional unit hyper-sphere. To do so, we generated a 9-dimensional vector with independent standard normal distributed elements for each sample and then normalized the vector to unit length.

